# Identification of cross-reactive vaccine antigen candidates in Gram-positive ESKAPE pathogens through subtractive proteome analysis using opsonic sera

**DOI:** 10.1101/2025.02.06.636871

**Authors:** Océane Sadones, Eliza Kramarska, Maite Sainz-Mejías, Rita Berisio, Johannes Huebner, Siobhán McClean, Felipe Romero-Saavedra

**Affiliations:** Division of Pediatric Infectious Disease, Hauner Children’s Hospital, LMU, Munich, Germany; Institute of Biostructures and Bioimaging, Italian Research Council (CNR), Naples, Italy; School of Biomolecular and Biomedical Sciences and UCD Conway Institute of Biomedical and Biomolecular Research, University College Dublin, Belfield, Dublin 4 D04 V1W8, Ireland

**Keywords:** cross-reactivity, antigen, enterococci, *Staphylococcus aureus*, vaccines, ESKAPE pathogens, opsonophagocytic assay, AdcA

## Abstract

The Gram-positive pathogens of the ESKAPE group, *Enterococcus faecium,* and *Staphylococcus aureus*, are well-known to pose a serious risk to human health because of their high virulence and numerous drug resistances, making them a real concern in healthcare settings worldwide. To narrow down the list of previously identified promising protein vaccine candidates, a combination of several antigen discovery approaches was performed, in particular a “false positive analysis” of peptides generated by trypsin shaving with a subtractive proteome analysis. The final list of nine potential antigens included AdcA_au_, a protein performing the same function as AdcA_fm_, an already discovered antigen in enterococci. Bioinformatic analyses revealed that AdcA_au_ and AdcA_fm_ share a sequence identity of 41.2% and that the conserved regions had a high antigenicity. AdcA_au_ was selected for further investigation and the results reported in this manuscript demonstrate the opsonic properties of AdcA_au_-specific antibodies against the *S. aureus* strain MW2, as well as their cross-binding and cross-opsonic activity against several *S. aureus, E. faecium*, and *E. faecalis* strains. This study suggests that further investigation of cross-reactive activities is a valuable tool for discovering new antigens that cover more than one clinically relevant pathogen.

## Introduction

The bacteria *Enterococcus faecium, Staphylococcus aureus, Klebsiella pneumoniae, Acinetobacter baumannii, Pseudomonas aeruginosa, and Enterobacter spp* are clustered and known as the ESKAPE group which poses a serious worldwide threat. The bacteria in this list were categorized as being able to “escape” the antibiotics’ biocidal action. [1]. The prevalence of these resistances in bacterial organisms is recognized as a global problem; in 2019, 1.27 million fatalities were attributed to these species globally; by 2050, that number is expected to climb to 10 million [2,3]. Two species of the ESKAPE group, *E. faecium,* and *S. aureus*, are the most relevant Gram-positive bacteria on the World Health Organization’s (WHO) priority list of antibiotic-resistant bacteria. They are ranked as “Priority 2: High” based on several factors, including but not limited to treatability, mortality, healthcare burden, and emerging resistance [4].

*Staphylococcus aureus* and enterococci are components of the commensal microbiota in healthy people [5]. When the microflora becomes imbalanced, however, these opportunistic organisms thrive and can cause potentially fatal illnesses such as bacteremia or endocarditis [6–9]. At this point, the length of hospital stay, the expense of therapy, and—above all—the mortality rate all increase substantially [10]. Antibiotic treatment is frequently unsuccessful, usually selecting resistant bacteria over sensitive bacteria in the case of common vancomycin-resistant enterococci (VRE) or methicillin-resistant *S. aureus* (MRSA) [11]. Together, VRE and MRSA account for 45% of deaths attributed to infections by antimicrobial-resistant pathogens in the United States in 2017 [12,13]. These infectious agents not only proliferate in clinical settings and long-term care units, where they are to blame for a significant portion of nosocomial infections but they are also isolated from environmental reservoirs, posing a threat to universal health care through community-acquired diseases [14].

While the discovery of antibiotics is still regarded as one of the most significant advances in medical research, antibiotic therapies are often unsuccessful in combating VRE and MRSA [15]. Another significant scientific advancement that offers great promise is vaccination. The use of antibiotics, the expense of treatment, and the length of hospital stay can all be reduced by preventing bacterial infections [10]. Attempts to develop a novel staphylococcal or enterococcal vaccine resulted in the identification of several possible vaccine antigen candidates, but no effective and commercially viable formulation has been achieved so far [16–18].

The current methods for antigen identification often lead to a large number of candidates. Testing the antigenic potential for extensive lists of proteins is time-consuming, labor-intensive, and expensive. To narrow down as much as possible the list of novel protein antigen candidates in *S. aureus*, several techniques were combined to reduce the number of identified proteins to only promising candidates that are targeted by opsonic antibodies. The techniques were already proven to be effective, either with the tested organism or in other bacterial species. Specifically, the methods combined a false-positive analysis on trypsin-shaved protein extracts with subtractive proteome analysis (SUPRA). Briefly, the “false-positive” strategy included the analysis of peptides generated by trypsin shaving controlled with peptides obtained using the same steps but incubated without the enzyme [19]. SUPRA was performed on protein extracts generated through lysostaphin digestion of the cell wall, sodium dodecyl sulfate (SDS) boiling, and sonication. Out of the nine identified candidates, one protein had a similar function with an already described antigen in *E. faecium*. As bioinformatic analysis showed a great sequence identity between the two proteins: AdcA_fm_ (AdcA from *E. faecium*) and AdcA_au_ (from *S. aureus*), the ATP-binding cassette (ABC) transporter was chosen for further investigation as a potential antigen. An immunological analysis of polyclonal sera generated in rabbits against the recombinantly produced proteins was used to investigate the potential efficacy of AdcA_au_ against both Gram-positive ESKAPE pathogens.

## Methods

The experiments described in this study were conducted from July 2020 to December 2023. This timeframe ensures the study’s epidemiological relevance and provides context for the data presented.”

### Bacterial strains and culture conditions

Table 1 presents the bacterial strains used in this study. The medium Brain Heart Infusion was used to grow *S. aureus* on plates at 37°C or in liquid media at 37°C with agitation. Enterococci were grown in Tryptic Soy Agar or Broth at 37°C without agitation. Proteins were produced using *Escherichia coli* M15 grown in Luria Bertani (LB) at 37°C with agitation. The bacterium harbored pRep4 and was therefore cultivated with LB containing kanamycin at 25 μg/ml. Ampicillin at 100 μg/ml was added after transformation with pQE30.

**Table 1:** Bacterial strains used for this study.

| Strains | Description | Assessed for research purposes on (dd/mm/yyyy): | Reference |
| --- | --- | --- | --- |
| *Staphylococcus aureus* |  |  |  |
| *S. aureus* MW2 | CA-MRSA isolate from the USA (North Dakota) | 15/12/1999 | [20] |
| *S. aureus* Reynold | Clinical isolate | 02/04/1997 | [21] |
| **Enterococci** |  |  |  |
| *E. faecium* 11236/1 | VRE isolated from a patient in Germany (Munich) | 22/08/2017 | [22] |
| *E. faecalis* 12030 | Isolated from a patient in the USA (Cleveland) | 18/01/1997 | [23] |
| *E. faecalis* Type 2 | Isolated from a patient in Japan (Sapporo) | 27/08/1991 | [24] |
| *Escherichia coli* |  |  |  |
| *E. coli* M15/pQE30AdcA<sub>fm</sub> | M15 harboring pRep4 and pQE30AdcA<sub>fm</sub> plasmids |  | [17] |
| *E. coli* M15/pQE30AdcA<sub>au</sub> | M15 harboring pRep4 and pQE30AdcA<sub>au</sub> plasmids |  | This study |

### Protein extraction by lysostaphin digestion

Protein extraction by lysostaphin digestion of the surface-exposed proteins was performed as described elsewhere [25]. *S. aureus* MW2 and MN8 were grown until the mid-exponential phase, the bacterial culture was harvested by centrifugation, and the pellet was washed twice in cold PBS (Phosphate Buffered Saline), followed by one wash with cold digestion buffer (PBS, 30% sucrose). Bacteria were resuspended in 2 ml of cold digestion buffer containing 200 μg of lysostaphin and incubated for one hour at 37°C under light agitation. Supernatant-containing protein was obtained by centrifugation at 2500 x g for 15 minutes at 4°C and clarified potential residues by centrifugation at top speed for two minutes and filtration with 0.22 μM filters.

### Protein extraction by SDS boiling

Preparation of protein extracts by SDS boiling was performed previously described by Nandakumar et al [26]. Briefly, bacteria were grown until OD_600nm_ reached 0.6. Cultures were harvested and washed twice with cold PBS. The pellet was resuspended in extraction buffer (PBS pH 8.0 + 2% SDS) at 10 μl per milligram of wet pellet. The suspension was heated at 95°C for three minutes and then placed back on ice. The proteins were isolated from the cells by centrifugation at 5,000 x g for 15 minutes at 4°C. The supernatant was centrifuged at top speed and run through a 0.22 μM filter to clarify potential residues.

### Protein extraction by sonication

The total protein extractions by sonication were performed using a protocol already described [27]. Bacterial cultures were grown to the mid-exponential phase and harvested by centrifugation. The pellet obtained was washed twice in cold PBS and resuspended in 4ml of cold PBS. The suspension was sonicated at 30% amplitude (750-watt Ultrasonic processor) for a duration of 20 minutes including on/off pulses of 9 seconds. Centrifugation at 10,000 x g for 20 minutes at 4°C was used to separate cell residues from extracted proteins. Further clarification was performed by filtration through a 0.22 μM filter.

### Trypsin shaving

Preparation of peptide extracts by trypsin shaving was performed as previously described [28]. Briefly, a bacterial culture was cultivated from OD_600nm_ 0.05 to 0.6. Cells were harvested and washed twice with cold PBS and once with cold digestion buffer (PBS, 30% sucrose). The pellet was resuspended in 2 ml of cold digestion buffer and the subsequent bacterial suspension was separated in two equal volumes. One volume was kept as it was and the other was used to resuspend 20 μg of trypsin. Both tubes were incubated for 15 minutes at 4°C with light agitation. Supernatants were collected by pelleting bacteria by centrifugation at 1,000 x g for 15 minutes at 4°C. Potential residues were removed by centrifugation of the supernatants at top speed and by filtration through a 0.22 μM filter.

### Depletion of pooled human sera

Sera isolated from eight healthy human donors were combined and are referred to as “HS” for human sera. Half of the pooled antibody volume was utilized to obtain sera depleted of *S. aureus*-specific antibodies, or “dHS” (depleted-HS). Bacteria were grown overnight, pelleted by centrifugation, and washed three times with PBS. After washing, the pellet was resuspended in 5ml of serum that had been diluted 1:10 in PBS and incubated with *S. aureus* MW2 or MN8 overnight at 4°C with gentle stirring. To get rid of any possible leftover bacteria, the suspension was centrifuged the next day, and the recovered supernatant was filtered.

### Immunodot blot

A series of dilutions of protein extractions – lysostaphin digestion, SDS boiling, and sonication -, ranging from 1:1 to 1:256, were performed and a 2 μl-drop of each dilution, and each protein extract, was placed on two independent PVDF membranes. The membranes were blocked overnight with a bovine serum albumin (BSA) solution. After washing with PBS containing 0.05% Tween20, the membranes were incubated for one hour with either HS or dHS, previously diluted at 1:2000. Membranes were washed again with the same washing buffer and incubated for one hour with a goat anti-human IgG coupled with HRP (horseradish peroxidase), diluted at 1:1000. After thorough washes, the membranes were detected using the PierceTM ECL Western kit, according to the manufacturer’s instructions.

### Subtractive proteome analysis

Protein samples were separated three times identically using 12% acrylamide SDS-PAGE. One gel was kept for staining with Coomassie blue and the two others were further analyzed by western immunoblotting. The gels were transferred onto Immobilon-P® polyvinylidene difluoride membranes. The membranes were activated by incubation in methanol for 30 seconds and 5 seconds in dH_2_O. After equilibration in transfer buffer (0.2 M Glycine, 25 mM Tris base, 20% methanol), the blotting was performed for 16-18h at 4°C and 30 mA (Bio-rad wet/tank blotting system) in wet conditions. The immunodetection was performed by incubation of the membranes with blocking buffer (5% BSA, 3% milk powder in PBS) overnight at 4°C to avoid unspecific binding. After careful washes, the membranes were incubated for 4 hours at 4°C with either HS or dHS for attachment to the recognized proteins. Membranes were then washed again and incubated for one hour at 4°C with a goat anti-human IgG coupled with HRP. After washing, membranes were detected by chemiluminescence on a Vilber Fusion Fx (Vilber, France) imaging system using the PierceTM ECL Western kit, according to the manufacturer’s instructions. The bands revealed by chemiluminescence were compared between the two membranes and bands identified with HS, but not with dHS were matched with the corresponding protein bands observed after destaining the associated gel. Those bands were excised using a clean scalpel, incubated in 100 mM ammonium bicarbonate (AB)/acetonitrile (ACN) (1:1) for 30 min-incubation, and occasionally vortexed for destining. The resulting destined gels pieces were incubated in neat ACN until they shrank and then incubated in trypsin buffer (13 ng/μL trypsin in 10 mM AB containing 10% ACN) for two hours on ice. 100 mM AB were added and the mixture was incubated overnight at 37°C to digest the proteins within the gel into peptides. The remaining pieces were then incubated with extraction buffer (5% formic acid/ACN (1:2)) at 37°C for 15 minutes to extract the digested peptides from the gel pieces. To exchange the buffer containing the digested peptides, supernatants were transferred to new tubes, dried in a vacuum, and redissolved in 0.1% trifluoroacetic acid. To properly get a solution and no remaining pieces of dried peptides, the solution was sonicated for five minutes and centrifuged for 10 min at 10,000 rpm, then finally dried in a vacuum Eppendorf Concentrator 5301.

### Mass spectrometry analyses

Protein extracts obtained by trypsin shaving and peptides obtained by band excision and in-gel digestion were analyzed by MALDI-TOF (Matrix Assisted Laser Desorption Ionization - Time of Flight) to determine the identity of proteins in each sample. For trypsin shaving, the enrichment of proteins in the treated samples was determined by applying different types of calculation for each protein identity found: difference in the number of peptides, in the percentage of peptides, in intensity, and label-free quantification intensity. Each type of calculation led to a list that was then ranked from highest to lowest difference and only the five top percent of each four lists were kept for further analysis. For samples excised from the SDS-PAGE, potential contaminants, proteins identified with a sequence coverage under 15% and, proteins having a molecular weight variation over 20% compared to the excised band, were removed from the tables giving the protein identities found in each band.

The frequency of each candidate’s identification was analysed and proteins discovered in both bacterial strains and using both techniques (false-positive analyses and SUPRA) were kept. Each remaining candidate was examined to create a shortlist based on predetermined standards. First, proteins that have previously been investigated as *S. aureus* antigens were excluded from the list. Bioinformatics and bibliographic research methods were used to gather information on the retained proteins: the immunogenicity, allergenicity, and subcellular localization were examined to narrow down the list of proteins.

### Bioinformatics

Sequence alignments were performed with EMBOSS Needle [29]. The subcellular localization was obtained on the UniProt website (https://www.uniprot.org/) by accessing information for each candidate through their UniProt number. Further bioinformatic analyses were conducted using prediction tools: CELLO v.2.5 (http://cello.life.nctu.edu.tw/) [30] and Gpos-mPLoc (http://www.csbio.sjtu.edu.cn/bioinf/Gpos-multi/) [31]. The Vaxijen tool was used for antigenicity prediction, which uses algorithms based on principal amino acid properties of a protein sequence [32]. Allergenicity was assessed using AllergenFP v.1 [33].

### Production of polyclonal sera in rabbits

*E. coli* M15 harboring pRep4 and the genetically engineered pQE30AdcA_fm_ or pQE30AdcA_au_ were used to recombinantly produce the proteins as previously described by Romero-Saavedra et al. (2015) [17]. *E. coli* M15 containing pRep4 were transformed with pQE30 genetically engineered to contain the gene encoding AdcA_au_. The gene was previously amplified by PCR using the primers presented in Table 2 and the genomic DNA of *S. aureus* MW2 as a template. After selection for positive transformants, bacteria were grown at 37°C until the optical density at 600 nm reached 0.5 and induced for two hours with 0.5 mM of isopropyl β-D-1-thiogalactopyranoside. After harvesting and washing, the bacteria were lysed enzymatically and mechanically. The His-tagged protein was isolated by affinity chromatography with a nickel resin and desalted and concentrated by diafiltration with the Amicon Ultra-15 Centrifugal Filter Units of 10,000 MWCO.

**Table 2:**
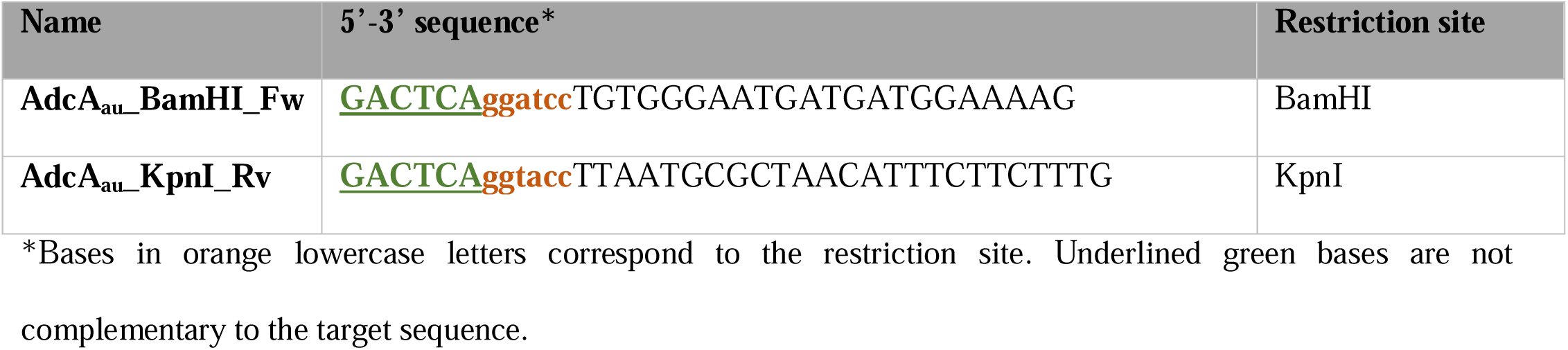
Primers used for this study.

Polyclonal serum was generated against AdcA_au_ by immunization of New Zealand rabbits, previously selected for a low pre-existing opsonic antibodies level. The immunization schedule included four subcutaneous injections, all containing protein and a proprietary adjuvant from Biogenes. After collecting pre-immune serum, rabbits were injected with 200 μg of recombinant protein at day 1. Following this first immunization, three boosts with 100 μg of recombinantly produced protein were injected at days 7, 14, and 42. At day 49, anti-protein serum was collected and heat-inactivated at 56°C for 30 minutes. The different rabbit sera used in this study are listed in Table 3.

**Table 3:** Sera used for this study.

| Sera | Description | Reference |
| --- | --- | --- |
| Pre-AdcA<sub>au</sub> | Pre-immune serum collected at day 0 from rabbits immunized with AdcA<sub>au</sub> | This study |
| Anti-AdcA<sub>au</sub> | Anti-protein serum collected at day 49 from rabbits immunized with AdcA<sub>au</sub> | This study |

### Immunodetection of specific antibodies by ELISA

The ELISA was carried out following the methodology previously reported by Romero-Saavedra et al. (2019) [34]. Recombinant proteins were utilized to coat Nunc-immuno MaxiSorp 96-well plates overnight at a concentration of 1 μg/ml. Following washing, plates were blocked using PBS-BSA (phosphate buffer saline with 3% bovine serum albumin) for one hour. Sera were adjusted at various concentrations between 60 and 15 μg/ml for cross-binding studies and between 1.8 and 0.45 μg/ml for specificity studies in PBS-BSA, added to the plate, and incubated for an hour. After washing, polyclonal alkaline phosphatase-conjugated anti-rabbit IgG produced in goat (Sigma; A3812-.25ML) was diluted to 1:1000 and added to the plates for a one-hour incubation. Ortho-nitrophenyl-b-D-galactopyranoside at 1 mg/ml was then added to the plates after the wells had been rinsed five times. After two hours, the absorbance was measured at 405 nm.

### Opsonophagocytic killing Assay

To perform the *in vitro* opsonophagocytic assay, antibodies, bacteria, complement, and white blood cells were prepared as previously described [23,35]. RPMIF (Roswell Park Memorial Institute 1640 medium with 15% fetal bovine serum) was used to dilute pre- and anti-protein sera to different dilutions, as well to dilute baby rabbit complement, to either 1:30 or 1:15, depending on the bacterial strains used, *S. aureus* or enterococci respectively. Baby rabbit complement was depleted from IgGs directed towards the pathogens by incubation with an excess amount of bacteria for one hour. Bacteria were cultivated until their OD_600nm_ reached 0.4 and then diluted in RPMIF following a prior buffer media exchange. White blood cells were freshly extracted from healthy volunteers and adjusted to a concentration of 18.10^6^ cells/ml. All four components were incubated at 37°C for 90 minutes and diluted and plated on agar plates. Colony-forming units were counted the day after and percentages of killing were calculated by comparing the numbers with or without white blood cells.

### Quantification and statistical analysis

The software GraphPad PRISM version 5.00 was used to statistically compare the data. For *in vitro* experiments, the unpaired two-tailed T-test with a 95% confidence interval was used. Bars and whiskers represent mean values ± standard error of the mean (SEM). NS, not significant (P > 0.05). *P ≤ 0.05, ** P ≤ 0.01, *** P ≤ 0.001.

### Ethics statements

Polyclonal sera raised in rabbits was performed by immunization by the company BioGenes GmbH in Berlin (Germany), following German guidelines and animal welfare regulations from the European Union, for housing, immunizing, and collecting serum samples. The study was approved by the Institutional Review Board NIH/OLAW (identifier F16-00178 (A5755-01), Approved 29 January 2024).

Human sera were collected from healthy donors as described in the study protocol approved by the Ethics committee at the LMU Munich, on June 1^st^, 2022 (Project number 22-0263). Consent was given by writing. Clinical strains were obtained from the collection of a bacteriological laboratory of the hospital and are therefore not associated with patients.

## Results

### Novel experimental design for antigen discovery

The experimental design followed is described in Fig 1. Several protein extractions were performed (Fig 1A) and treated in different ways. A false-positive analysis of trypsin shaving extractions was combined to a novel version of the previously described subtractive proteome analysis, for which three different protein extractions were carried out: either by lysostaphin digestion of the peptidoglycan in hypotonic solution; by boiling in an SDS-containing buffer; or using sonication. To conduct the subtractive immunoblotting, antibodies from healthy human donors was used after being pooled together (HS) and depleted of *S. aureus*-specific antibodies by incubation with the whole bacterium (dHS) (Fig 1B). The protein extracts were separated by SDS-PAGE (sodium dodecyl sulfate-polyacralamide gel electrophoresis) concurrently, in triplicate. Two of the replicates were blotted onto a polyvinylidene difluoride (PVDF) membrane and detected with either the HS or dHS (Fig 1C). Bands identified with the non-depleted serum but not with the depleted serum were matched to the bands on the gel and excised. After gel digestion, the samples were analysed by MS to identify the proteins present in the excised bands, likely to be proteins that are the targets of opsonic antibodies, and therefore promising vaccine candidates.

**Fig 1:**
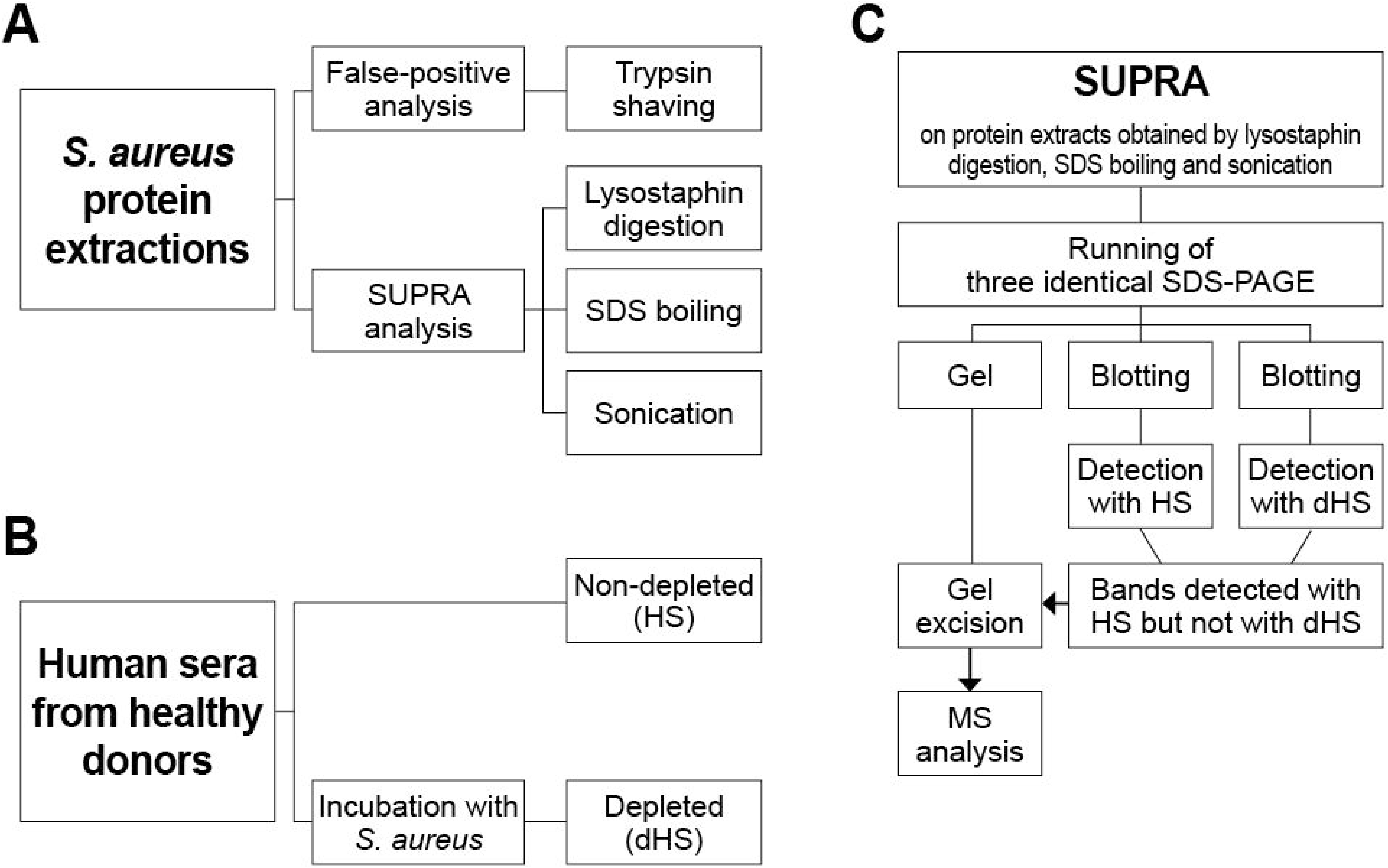
Experimental design for antigen discovery. (A) Preparation of protein extractions from *S. aureus* MW2 and *S. aureus* MN8. (B) Acquisition of antibody input for SUPRA technique. (C) SUPRA analysis on different protein extracts using HS or dHS.

### Human sera depletion of *S. aureus*-specific antibodies

Human sera from healthy donors were depleted of *S. aureus*-specific antibodies as shown in Fig 2A. The efficiency of the depletion was examined by immunodotblot (Fig 2B). For *S. aureus* MW2, the intensity of the spots in the membrane incubated with HS is higher than the membrane incubated with dHS, meaning that protein recognition by dHS was lower than that of HS, especially for the protein extract obtained by sonication. Indeed, while spots were detected until a dilution of 1:128 with HS, none was detectable with dHS, even without diluting the sample. For SDS boiling, protein spots are still detected at 1:256 with HS, while not anymore from 1:32 with dHS. For lysostaphin-obtained protein extracts, detection still occurs at 1:128 for HS, but only until 1:16 with dHS. These results show that while the depletion of *S. aureus*-specific antibodies is not entirely achieved with the strain MW2, the protocol still greatly reduced the number of antibodies able to bind to the protein extracts. In the case of *S. aureus* MN8, each protein extract can be detected by HS, up to 1:64 for the lysostaphin-mediated extraction, 1:16 for SDS boiling, and 1:32 for sonication. However, no detection was possible using dHS, meaning that the depletion was effective.

**Fig 2:**
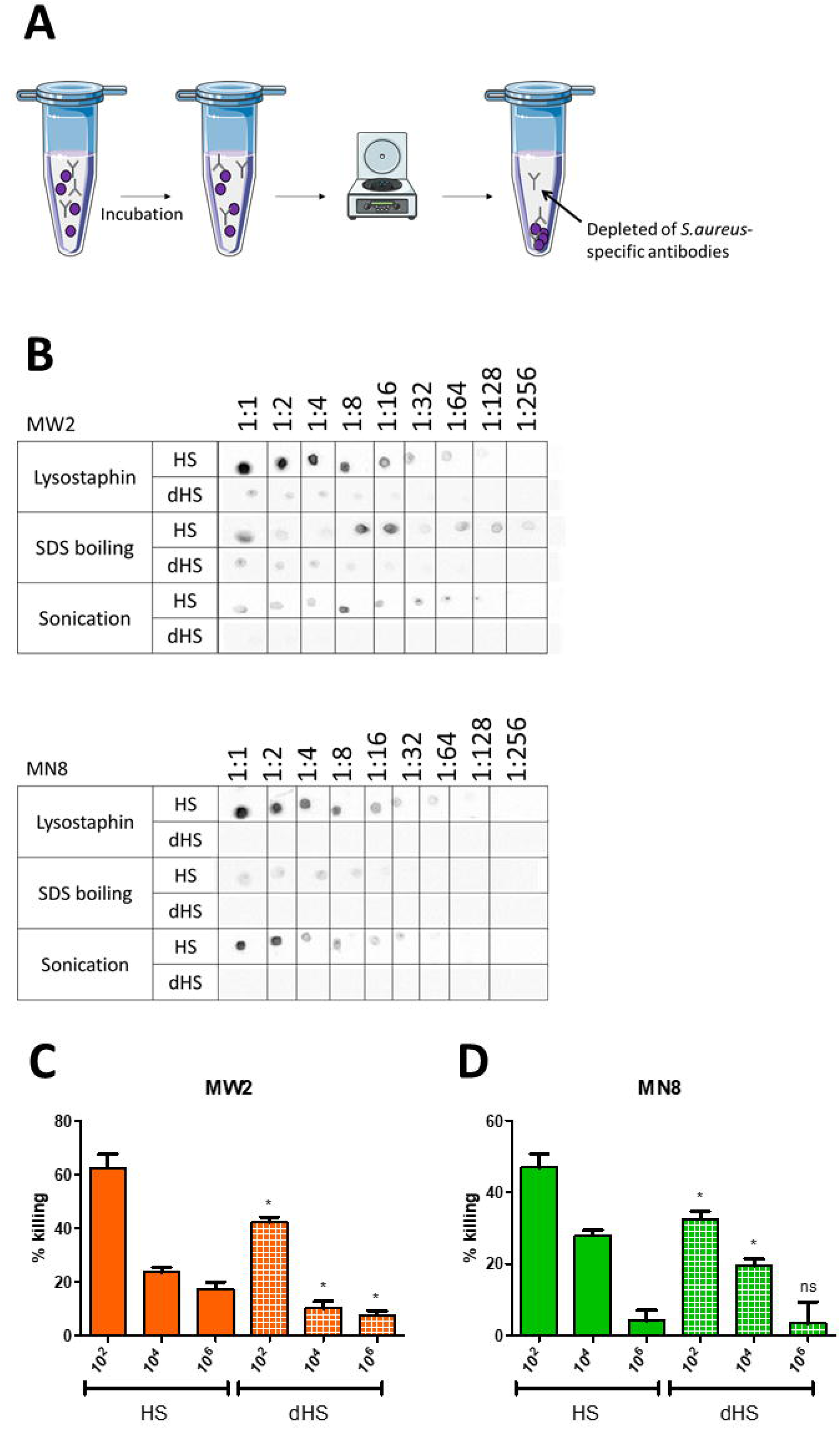
Human sera depletion of *S. aureus* specific antibodies. (A) Depletion protocol. Human sera were placed with *S. aureus* MW2 or MN8 (purple spheres) and incubated overnight. (B) Comparison of detection intensity of the different protein extractions with HS and dHS using immunodot blot. The different protein extractions were diluted with a factor of 2 until 1:256. A drop of each dilution was placed on a PVDF membrane. The membrane was blocked with BSA and further incubated with HS or dHS, followed by incubation with a conjugated anti-human IgG. (C, D) OPA against *S. aureus* MW2 (orange) or MN8 (green) using HS (plain bars) or dHS (white squares). Values from the same dilution of HS and dHS were compared statistically using an unpaired two-tailed T-test with a 95% confidence interval. Bars and whiskers represent mean values ± standard error of the mean (SEM). NS, not significant (P > 0.05), *P ≤ 0.05.

To further investigate the success of the depletion, but also to check for the *in vitro* efficacy of the antibody’s input for the SUPRA technique, opsonophagocytic assays (OPA) were performed. Human sera and depleted-human sera were used against *S. aureus* MW2 (Fig 2C) and MN8 (Fig 2D). The results show a dose-dependent killing using each tested serum against both strains. Killing with dHS is significantly lower when compared to the same dilutions of HS, with reduced opsonic killing by approximately 16.5% for both strains. The depletion protocol was successful in reducing the opsonic killing mediated by the antibodies present in the human sera.

### Identification of potential protein targets

Following the successful depletion of pooled human sera, the SUPRA was performed to identify potential protein targets. The pictures of gels and associated detection membranes are presented in Fig 3. As expected given the previous immunodot blot results (Fig 2B), blotted membranes detected with dHS showed lower signal intensity when compared to HS. In the case of *S. aureus* MW2, comparing both membranes led to the detection of two bands showing a signal with HS but not dHS. Using a similar comparison, seven positive hits for MN8 were observed. Using the molecular maker and the pattern observed in the membrane incubated with HS (which shows more bands), interesting bands in the gels were identified and are highlighted in orange. Each band was carefully excised from the gels and treated for MS analysis.

**Fig 3:**
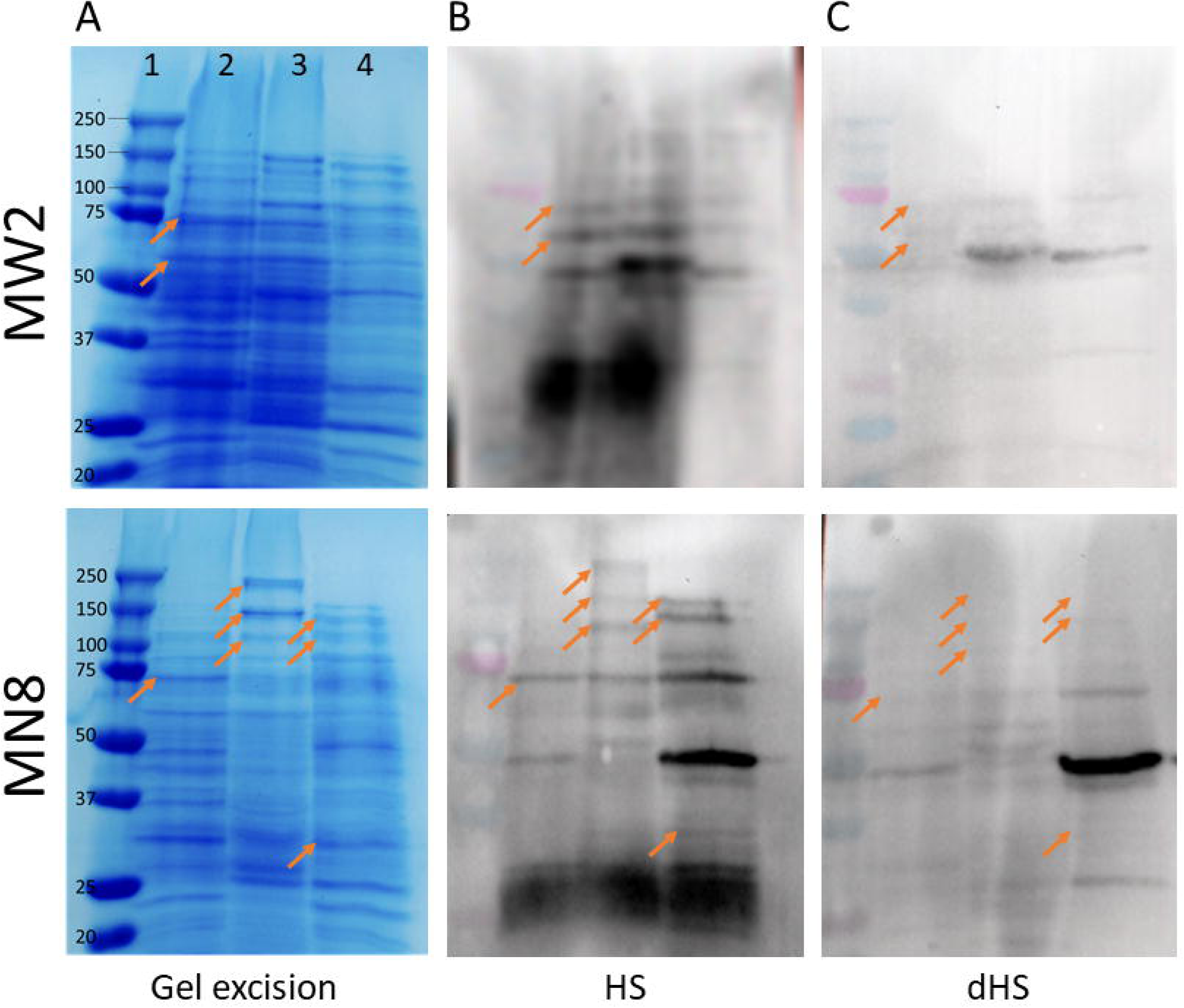
Identification of potential protein targets by using the SUPRA technique. (A) Different protein extracts run on a SDS-PAGE and associated excised spots. Line 1, molecular marker. Line 2, sonication. Line 3, lysostaphin. Line 4, SDS boiling. (B) Blotted gel on membrane detected with HS. (C) Blotted gel on membrane detected with dHS. Arrows show the bands considered for excision.

### Novel analysis pipeline from MS results

Proteins in the trypsin shaving samples and their associated controls, as well as proteins excised from the gels, were identified by MS using MALDI-TOF. Surface-exposed proteins were identified following a false-positive analysis of samples obtained by trypsin shaving: bacterial cells were incubated with (TS) or without any enzyme (cTS). Aiming to show the enrichment of proteins due to the trypsin activity, four calculations to highlight the differences between TS and cTS were performed as described in the method’s paragraph “Mass spectrometry analyses”. The list of proteins identified by MS from samples obtained by gel excision of interesting bands was trimmed following instructions given in the Methods section.

Pooling the results of trypsin shaving and the SUPRA technique gave a list of 40 potential candidates, identified with both methods. From this list, several characteristics were evaluated to further shortlist the potential antigens (see Methods section). As a result, nine promising antigen candidates that have not yet been studied were identified and are presented in Table 4.

**Table 4:**
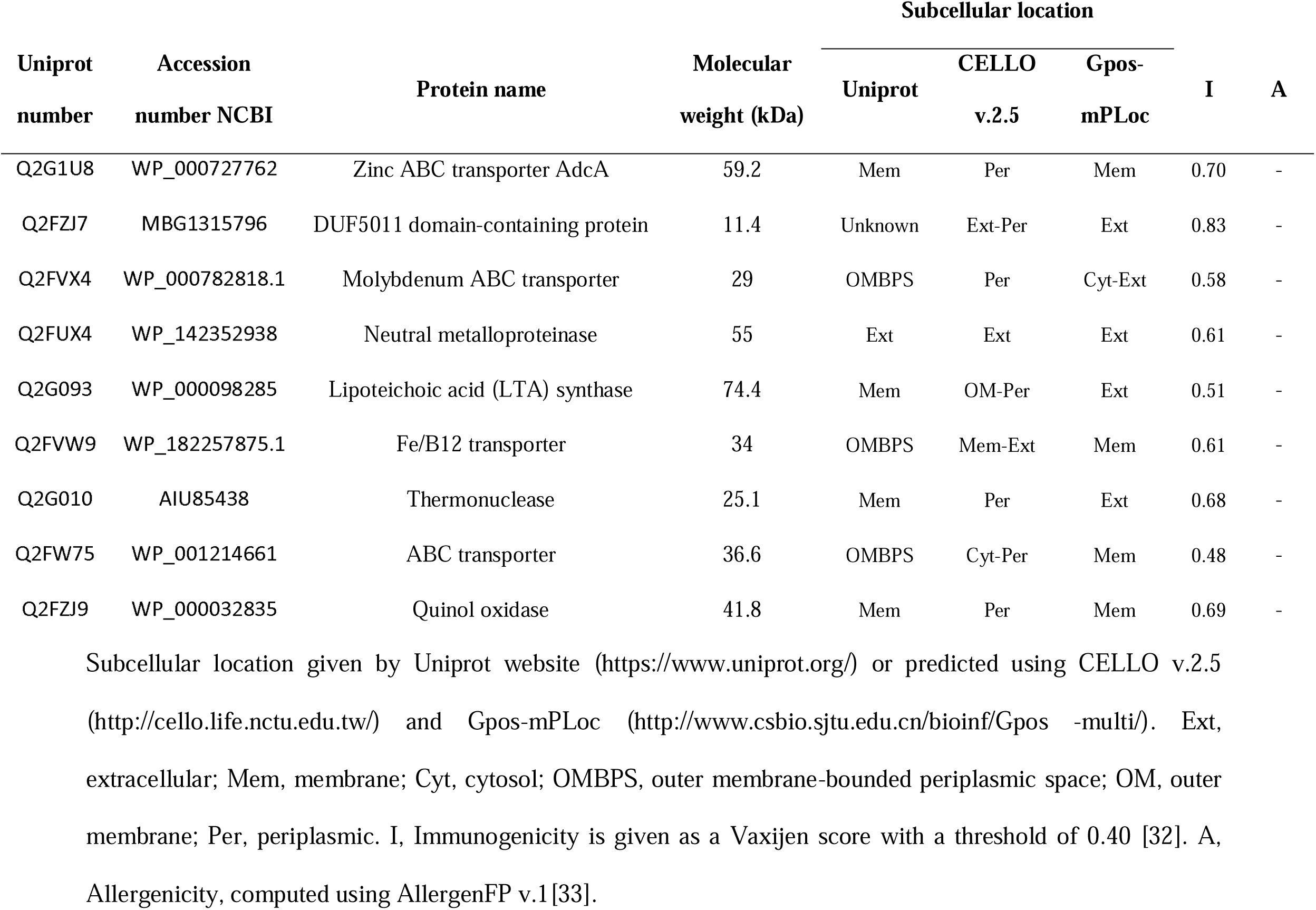
Summary of the candidates identified and their corresponding criteria.

### Sequence homology of AdcA_au_ with to AdcA_fm_

The zinc ABC transporter AdcA identified by combination of antigen discovery techniques is a ZinT domain-containing protein that is involved in zinc uptake and a protein with a similar function (AdcA_fm_) was already described as a protective antigen against both *E. faecium* and *E. faecalis* [34].Taking that into account, a pairwise alignment was performed to check if the similarity in function was translated to the sequence level (Fig 4). The overall pairwise sequence alignment of AdcA_au_ and AdcA_fm_, computed using EMBOSS Needle, is 41.2%. Also, many strongly conserved regions (>60% sequence identity) are highlighted, both in the ZinT and the ZnuA domains. The Vaxijen tool was used to assess the predicted antigenicity of the full-length proteins and the strongly conserved regions. Consistently, both proteins showed a high immunogenicity score, and conserved regions were predicted to be antigenic as well (Table 5).

**Fig 4:**
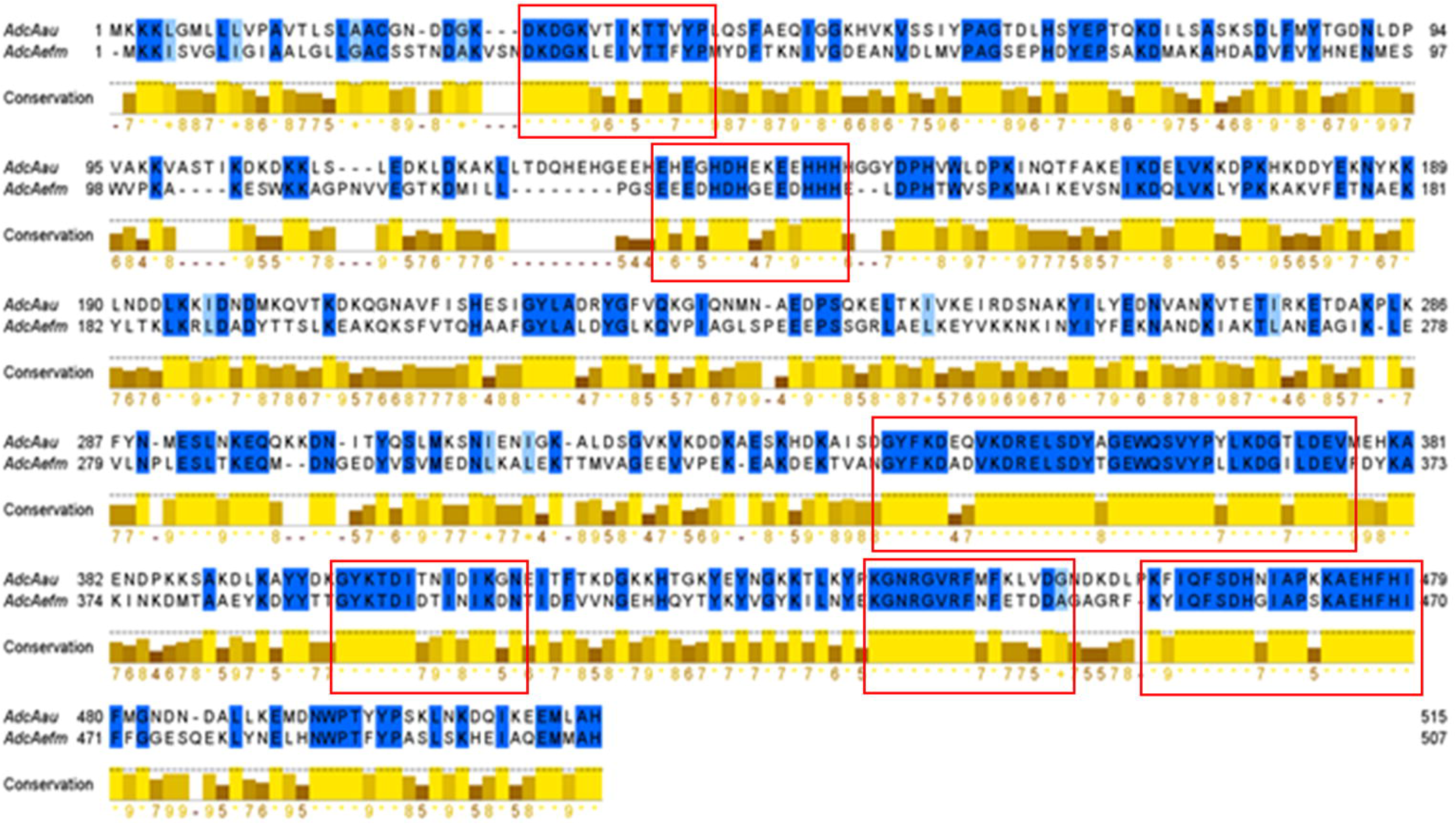
Pairwise alignment of AdcA_au_ and AdcA_fm_ using EMBOSS Needle. Red boxes mark regions of minimum length 14 residues presenting a sequence identity above 60%.

**Table 5.** Predicted immunogenicity of most conserved regions of AdcA_au_ and AdcA_fm_.

| Protein | Sequence | Vaxijen score | Sequence identity |
| --- | --- | --- | --- |
| AdcA<sub>au</sub> | 28-DKDGKVTIKTTVYP-41 | 1.10 | 71.4% |
| AdcA<sub>fm</sub> | 31-DKDGKLEIVTTFYP-44 | 0.87 |  |
| AdcA<sub>au</sub> | 133-EHEGHDHEKEEHHH-146 | 1.70 | 64.3% |
| AdcA<sub>fm</sub> | 127-EEEDHDHGEEDHHH-140 | 1.57 |  |
| AdcA<sub>au</sub> | 342-GYFKDEQVKDRELSDYAGEWQSVYPYLKDGTLDEV-376 | 0.43 | 85.7% |
| AdcA<sub>fm</sub> | 334-GYFKDADVKDRELSDYTGEWQSVYPLLKDGILDEV-368 | 0.42 |  |
| AdcA<sub>au</sub> | 399-GYKTDITNIDIKGN-412 | 1.73 | 71.4% |
| AdcA<sub>fm</sub> | 391-GYKTDIDTINIKDN-404 | 1.58 |  |
| AdcA<sub>au</sub> | 439-KGNRGVRFMFKLVDG-453 | 1.38 | 66.7% |
| AdcA<sub>fm</sub> | 431-KGNRGVRFNFETDDA-445 | 1.67 |  |
| AdcA<sub>au</sub> | 460- KFIQFSDHNIAPKKAEHFHI-479 | 0.68 | 85% |
| AdcA<sub>fm</sub> | 451-KYIQFSDHGIAPSKAEHFHI-470 | 0.58 |  |

### Immunization of rabbits with AdcA_au_ leads to the production of specific and cross-binding antibodies

To study the potential of AdcA_au_ as an antigen, polyclonal serum was produced in rabbits against the candidate. The protein was recombinantly produced and used for rabbit immunization. Pre-immune serum (pre-AdcA_au_) was isolated at day 0, before immunization with the recombinant protein. After several injections, anti-protein serum (anti-AdcA_au_) was collected on day 49. Both rabbit sera were used in ELISA (enzyme-linked immunosorbent assay) to determine whether the immunization led to the production of AdcA_au_-specific antibodies. Signals obtained with anti-AdcA_au_ are significantly higher when compared to the signals reflecting the binding of pre-AdcA_au_ (Fig 5A). Also, the absorbances measured when adding anti-AdcA_au_ to the plates decreased in a dose-dependent manner. These data demonstrate that the immunization with AdcA_au_ led to the production of antibodies specifically binding to AdcA_au_. Additionally, experiments to investigate the cross-binding capabilities of the antibodies raised against the staphylococcal protein were performed. The absorbance reading indicates a significant increase in AdcA_fm_-binding antibodies in anti-AdcA_au_ when compared to the pre-immune serum (Fig 5B). Moreover, the signals acquired in the different dilutions decrease with the concentration of antibodies, showing a dose-dependent effect. These results confirmed that AdcA_au_ and AdcA_fm_ share one or several epitopes, the previously presented hypothesis based on bioinformatics predictions. It also suggests that AdcA_au_ might have potential as a cross-reactive candidate against the Gram-positive pathogens of the ESKAPE group.

**Fig 5:**
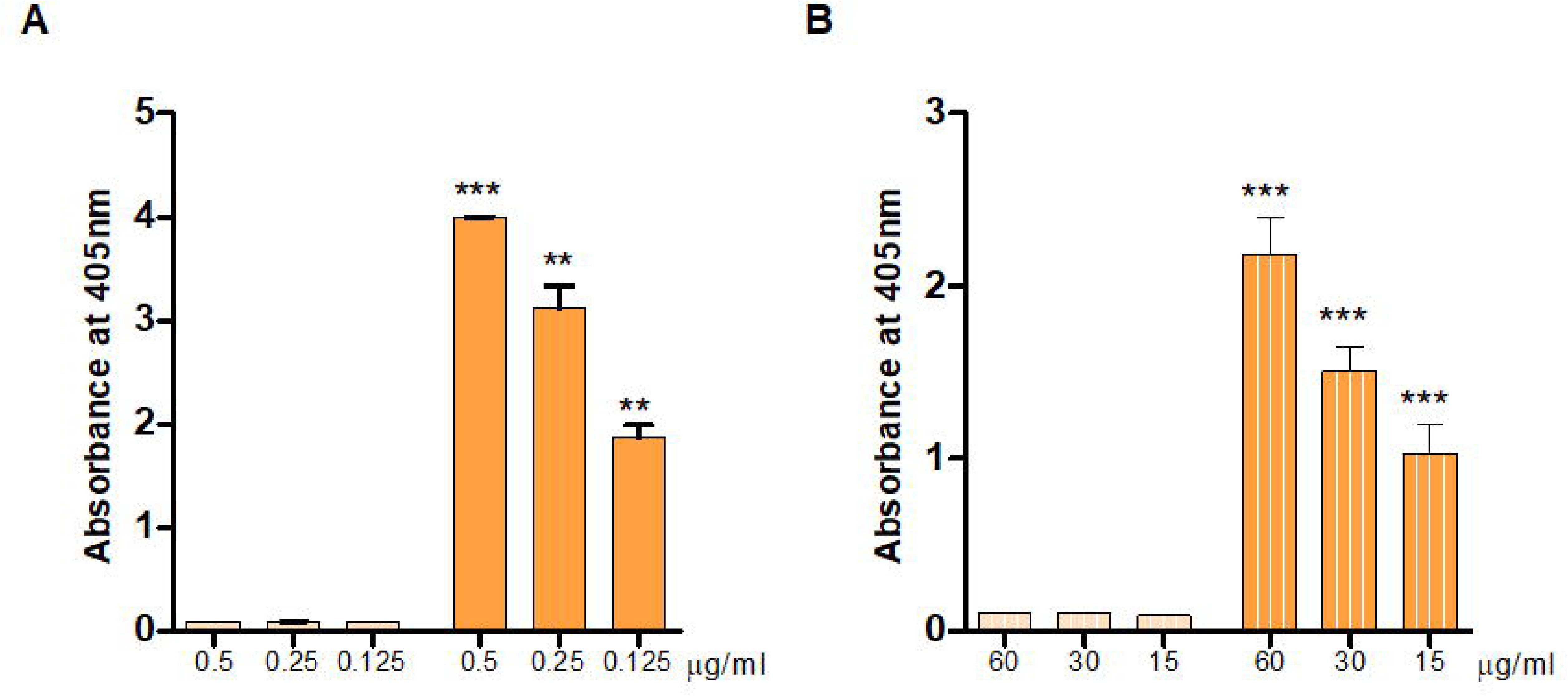
Immunoreactivity towards recombinantly produced proteins. (A) The ELISA technique was used to assess the binding of pre- (light orange) and anti-AdcA_au_ (dark orange) to the staphylococcal protein AdcA_au_. The His-tagged protein was used to coat Nunc-Immuno MaxiSorp 96-well plates. The rabbit sera were added at various concentrations to the wells. Results are shown as the absorbance obtained after detection by a conjugated secondary antibody and incubation with the substrate. (B) Using the same protocol, but coating the plates with a recombinantly produced AdcA_fm_, the cross-binding capabilities of pre- (light orange – stripes) and anti-AdcA_au_ (dark orange – stripes) were evaluated. An unpaired two-tailed T-test with a 95% confidence interval was used to statistically compare the absorbance values obtained for the same dilutions. Bars and whiskers represent mean values ± standard error of the mean (SEM). ** P ≤ 0.01, *** P ≤ 0.001.

### Polyclonal antibodies raised against AdcA_au_ mediate specific opsonic killing of *S. aureus* MW2

To investigate the potential of AdcA_au_ as an antigen, the polyclonal sera previously obtained by immunization of rabbits with the protein was tested. Pre- and anti-AdcA_au_ were used in OPA against *S. aureus* MW2 (Fig 6A). The killing mediated by opsonic antibodies was significantly higher with the anti-protein serum when compared to the pre-AdcA_au_. The increasing dilutions of anti-AdcA_au_ presented with a lower killing, with 1:320 being twice lower than 1:40 showing a dose-dependent effect of the polyclonal serum. Those results suggest a great *in vitro* efficacy of antibodies raised against the staphylococcal protein AdcA_au_.

**Fig 6:**
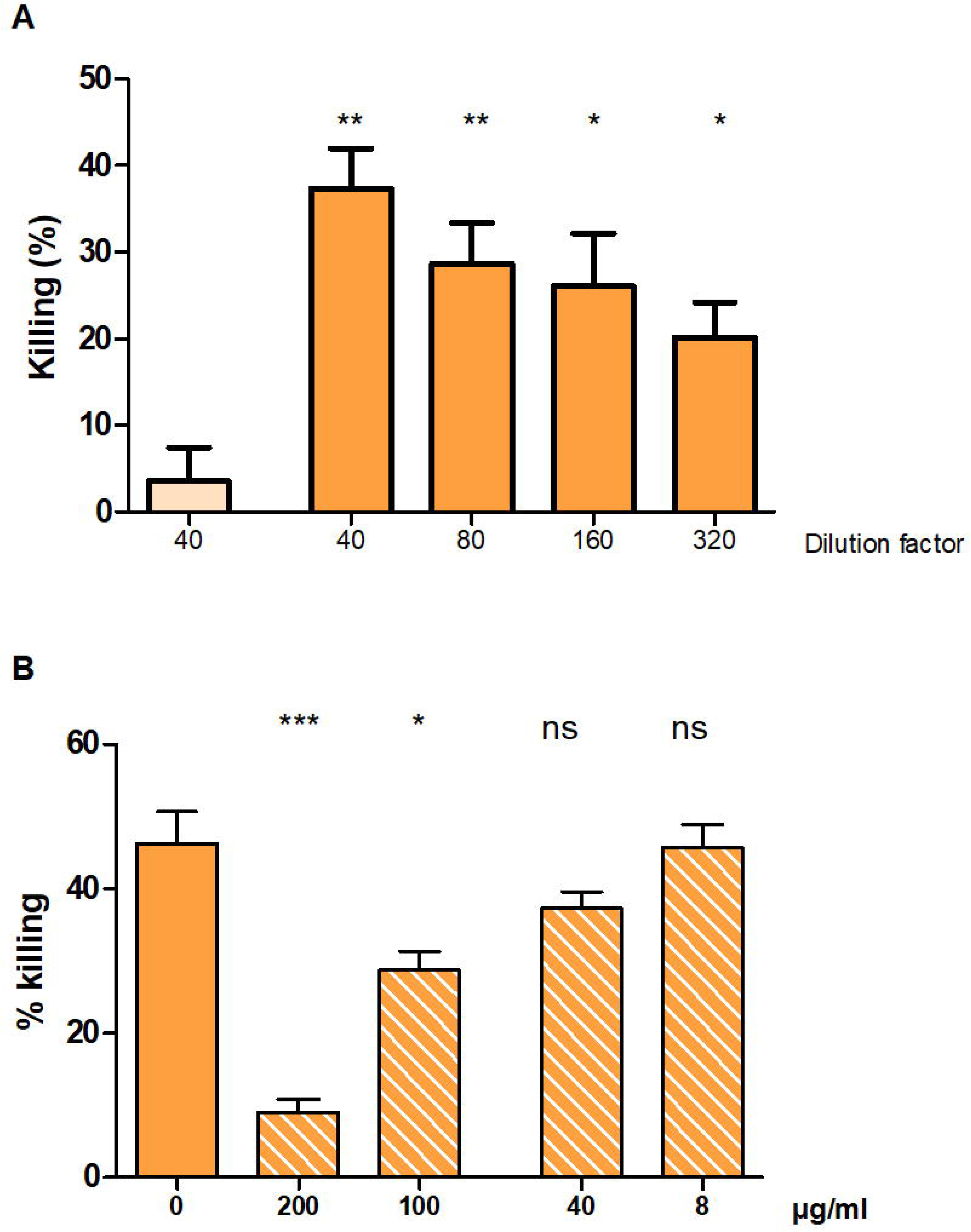
Opsonophagocytic killing assays against Gram-positive ESKAPE pathogens. (A) The opsonic killing of pre- (light orange) and anti-AdcA_au_ (dark orange) was assessed by OPA against *S. aureus* MW2. Values obtained for pre- and anti-AdcA_au_, at the same dilution, were statistically compared. (B) Inhibition of the killing mediated by anti-AdcA_au_ was performed using the recombinantly produced AdcA_au_ at different concentrations ranging from 8 to 200 μg/ml. The mediated killings observed for each antigen concentration were statistically compared to the mediated killings observed without protein. Values were compared using an unpaired two-tailed T-test with a 95% confidence interval. Bars and whiskers represent mean values ± SEM. NS, not significant (P > 0.05), *P ≤ 0.05, ** P ≤ 0.01, *** P ≤ 0.001

By using opsonophagocytic inhibition assay against *S. aureus* MW2, the specificity of the observed killing was evaluated (Fig 6B). The recombinantly produced AdcA_au_ was incubated with polyclonal sera for overnight at doses varying between 200 and 8 μg/ml. The inhibited sera were employed as an antibody source in OPA the following day. The findings indicate a dose-dependent reduction in killing caused by incubating the sera with the recombinant proteins. This shows that the mediated killing is uniquely caused by antibodies produced against AdcA_au_, as it may be prevented by incubation with the protein.

### Anti-AdcA_au_ contains antibodies mediating the killing of several *S. aureus* and enterococci

To test whether AdcA_au_ could also be used as an antigen covering more staphylococcal strains, both rabbit sera were tested against *S. aureus* Reynolds. Also, as shown by ELISA, antibodies raised against AdcA_au_ can bind to the enterococcal protein AdcA_fm_. Therefore, anti-AdcA_au_ was used to determine whether it could also mediate the killing of several enterococci: vancomycin-resistant *E. faecium* 11236/1, as well as *E. faecalis* 12030 and *E. faecalis* Type 2. The OPA results presented in Fig 7 show the opsonic killing mediated by pre-and anti-AdcA_au_. Values obtained with the anti-protein sera are significantly higher when compared to the pre-immune serum. Immunization of rabbits with AdcA_au_ led to the production of antibodies capable of mediating the opsonic killing of several strains of *S. aureus*, as well as enterococcal strains. Altogether, these results suggest that AdcA_au_ is a potential candidate antigen for vaccine formulation to prevent infections by both Gram-positive ESKAPE pathogens.

**Fig 7:**
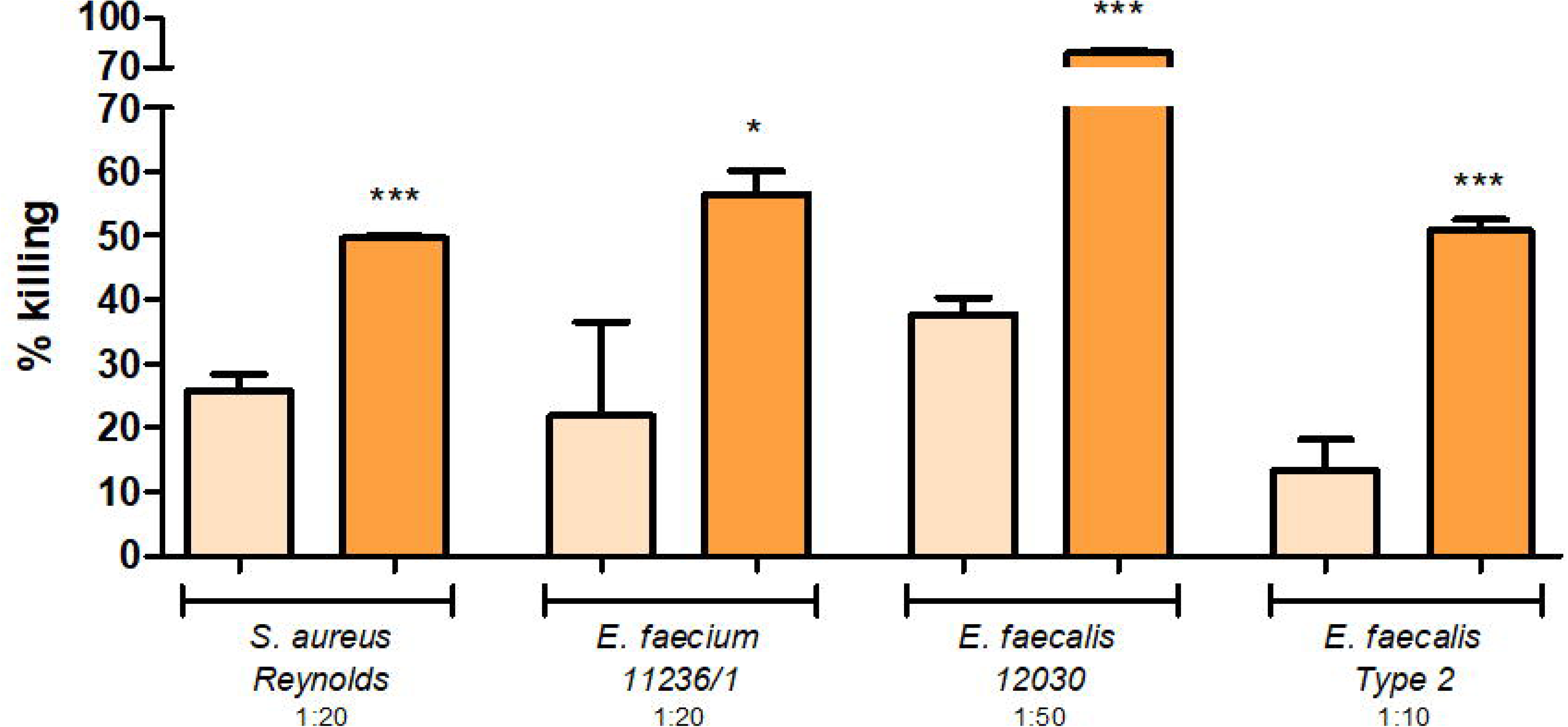
Cross-opsonic activity of anti-AdcA_au_. Several strains were tested to see the opsonic killing mediated by pre- (light orange), and anti-AdcA_au_ (dark orange). An unpaired two-tailed T-test with a 95% confidence interval was used to calculate the statistical difference between the values obtained with pre- and anti-AdcA_au_. * P ≤ 0.05, *** P ≤ 0.001.

## Discussion

Gram-positive ESKAPE pathogens pose a serious risk in the clinical setting [1,11,36]. In the United States between 2015 and 2021, enterococci and *S. aureus* were linked to 24% of all healthcare-associated infections [37]. Additionally, they are the most frequent organisms causing nosocomial bloodstream infections [36,38]. Effective vaccinations would be a huge benefit since they not only prevent serious diseases but also have been linked to lower antibiotic use, treatment costs, and shorter hospital stays [10,39,40]. Unfortunately, no vaccine has been successfully created to prevent infections by enterococci or *S. aureus*, despite several trials and a large number of identified antigens [15,16,22,34,41–45].

The purpose of this work was to identify novel *S. aureus* antigens and evaluate the efficacy of polyclonal sera generated against the discovered protein to ascertain its potential as antigens. Several previously published techniques were combined to reduce the number of potential candidates to a manageable quantity for further experimental investigation. The trypsin shaving technique consists of swelling up the bacteria by incubation in a hypotonic solution, leading to a better exposition of surface-exposed proteins that can be cleaved by the enzyme [28]. The peptides released in the supernatant were analysed by mass spectrometry (MS) and results were analysed following a so-called “false-positive analysis”, which consists in comparing the treated sample and control to reveal the enrichment of proteins in the supernatant due to trypsin digestion. This methodology aims to identify surface-exposed proteins, a common characteristic of vaccine candidates. This approach was combined with results obtained by protein extractions obtained by lysostaphin digestion of the peptidoglycan [25], SDS boiling [26], and sonication [27], using a subtractive proteome technique. SUPRA was performed using human sera from healthy donors and the same sera previously depleted for *S. aureus*-specific antibodies. The study yielded a list of nine candidates that were predicted to be surface-exposed [30,31], immunogenic [32], non-allergenic [33], and not already described as antigens against *S. aureus* in the literature [15]. From this list, AdcA_au_ was a homolog of a previously described bacterial antigen in enterococci: The ABC transporter substrate-binding lipoprotein from *E. faecium* (AdcA_fm_) [44]. The two proteins presented with highly conserved regions that were predicted to be immunogenic the potential of AdcA_au_ as an antigenic protein against both Gram-positive ESKAPE pathogens was investigated. ELISA experiments show that polyclonal sera targeting the staphylococcal protein can bind to both AdcA_au_ and AdcA_fm_. Using OPA, it was demonstrated that anti-AdcA_au_ can mediate the specific killing of several *S. aureus* strains and that the antibodies raised by immunization are cross-opsonic against various strains of both *E. faecium* and *E. faecalis* [46].

Proteins that act as part of a transport system can often be good potential vaccine antigens, especially due to their location at the cell surface. In the study reporting AdcA_fm_ as a good vaccine antigen candidate, Romero-Saavedra and colleagues also presented a list of surface-related proteins up-regulated in a mice peritonitis model [44]. While the antigenic potential of each protein listed was not investigated *in vitro*, they were considered as potential vaccine candidates against infections produced by this species. Among the 18 candidates in the list, three were part of a sugar transport system and two were involved in the transport of metals: the manganese ABC transporter substrate-binding lipoprotein PsaA_fm_ and the zinc ABC transporter substrate-binding lipoprotein AdcA_fm_. The two metal-binding proteins were investigated as vaccine antigens against enterococcal infections and showed promising results *in vitro* and *in vivo*. Interestingly, the protein MntC of *S. aureus* is also a manganese transporter that has been described as a vaccine antigen [47]. The antigen is part of the Olymvax vaccine formulation rFSAV currently studied in phase II of a clinical trial (clinical trial number NCT03966040) [48].

AdcA_fm_ is a substrate-binding protein in the ABC transport system and belongs to the metal-binding lipoproteins family. This protein regulates zinc (Zn) homeostasis in *E. faecium* [49]. Zn is a d-block transitional metal ion vital to all forms of life, has a structural and catalytic role in around 5% of bacterial proteins, where it participates in many biological processes such as carbon metabolism and DNA transcription regulation [50]. Bacteria acquire Zn via Zn-binding proteins, such as AdcA_fm_ and AdcA_au_, in order to survive in hostile environments and overcome the host-derived ion limitation [51,52]. Mutations in Zn-regulating genes consistently reduce virulence and enhance sensitivity to the host defense machinery in a variety of bacterial species [53–57]. Also, inactivation of the gene encoding the protein causes significant growth and survival defects as well as diminished resistance to ampicillin, bacitracin, and daptomycin [49]. Interestingly, the protein’s expression has been shown to increase during infection, and proteins with similar functions have already been investigated as virulence factors and/or potential vaccine antigens in other Gram-positive pathogens [53,58,59]. As Zn has a crucial role in bacteria, it is essential for bacteria to possess an ion uptake mechanism. These transport systems are found in many bacteria, including pathogens such as *Streptococcus pyogenes* and *Streptococcus pneumoniae*, where they play a role in the bacterial pathogenicity [60,61]. It is therefore not entirely surprising to observe a cross-reactive effect of antibodies targeting such proteins among different bacteria.

Altogether, the data demonstrate that the antigen discovery method used was successful in identifying a reasonable number of proteins as candidates. Also, the work shows that the consideration of proteins already described as antigens and exerting a crucial role within a bacterial cell should be further analysed for cross-reactive activity. It can also be noted that prevention of infections by several other streptococci may be possible, given the presence of a similar ZinT domain in those organisms [53,61]. While the main focus of this study was the study of AdcA_au,_ due to its similarity in function and sequence to AdcA_fm_, it is important to consider the investigation of the other eight candidates as potential antigens. The finding strongly supports the idea that pan-vaccinomic approaches could and should be based on the investigation of antigenic properties of proteins that play essential roles within the bacterium, as is the case with AdcA_au_ reported here.

## Conclusion

The study investigated and proved the potential of a newly identified antigen in *S. aureus*, AdcA_au_, which could be used in a vaccine formulation to prevent infections not only against *S. aureus* but also against the dangerous pathogens *E. faecium* and *E. faecalis*The experimental design successfully uncovered several promising vaccine candidates and could be applied to other bacteria in order to find new targets for vaccine design. Studying cross-reactive activity of antigenic proteins playing a crucial role for bacterial cells offers significant hope in the ongoing fight against antibiotic resistance and infection control caused by WHO-listed pathogens.

## Acknowledgements

O.S. wishes to acknowledge Mrs. Bernadette Freminet’s support, whose guidance led to the success of this research. The authors wish to thank Mrs. Corine Sadones for the help given to design the figures of this manuscript. Many thanks to Dr. Eugene Dillon for his help in analysing the mass spectrometry results. Mr Julien Hubert’s experimental support is also acknowledged.

## Author contributions

F. R.-S. and J. H. designed the research and reviewed the manuscript. O. S. performed the experiments and prepared the manuscript. E. K. and R. B. carried out the bioinformatic analysis. M.S.-M. and S.M. helped to design the proteomic experiments, as well as for the analysis of the results obtained. All authors read and approved the final manuscript.

## Supporting information

**S1_raw_images** file contains the uncropped images on gels end blots presented in this study.

