## Supporting information for "Identification of cross-reactive vaccine antigen candidates in Gram-positive ESKAPE pathogens through subtractive proteome analysis using opsonic sera"

#### Figure containing gels and blot images: Figure 3

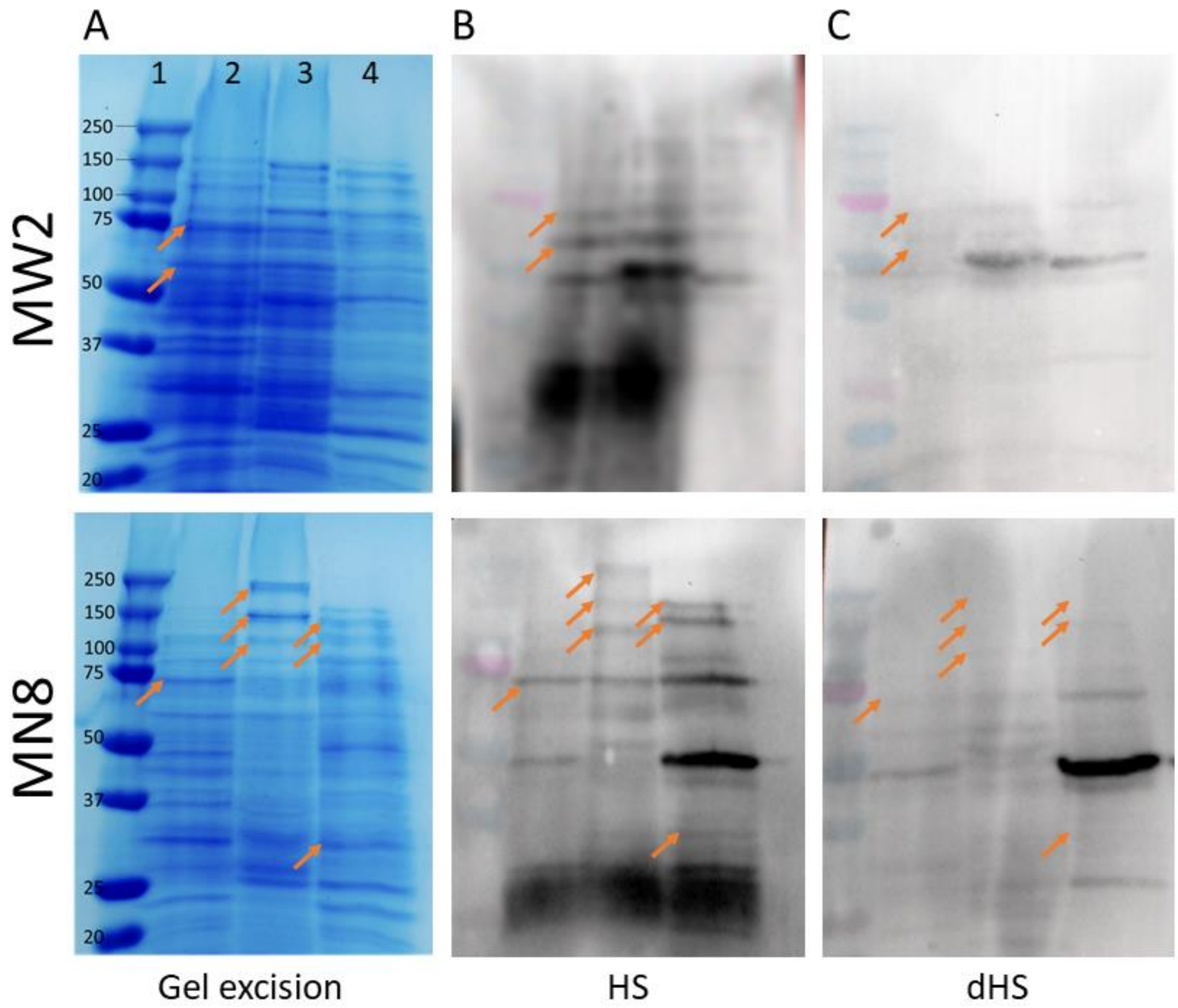


Fig 3: Identification of potential protein targets by using the SUPRA technique.

(A) Different protein extracts run on a SDS-PAGE and associated excised spots. Line 1, molecular marker. Line 2, sonication. Line 3, lysostaphin. Line 4, SDS boiling. (B) Blotted gel on membrane detected with HS. (C) Blotted gel on membrane detected with dHS. Arrows show the bands considered for excision.

### Raw images of gels and blots presented in Figure 3

#### Information

**Labelling of the raw images –** Each raw image is annotated with the numbers 1, 2, 3 and 4. 1 corresponds to the molecular marker, 2-4 indicate protein extractions obtained by sonication (2), lysostaphin digestion (3) or SDS boiling (4).

Molecular marker “Precision Plus Protein Dual Color Standards, #1610374”

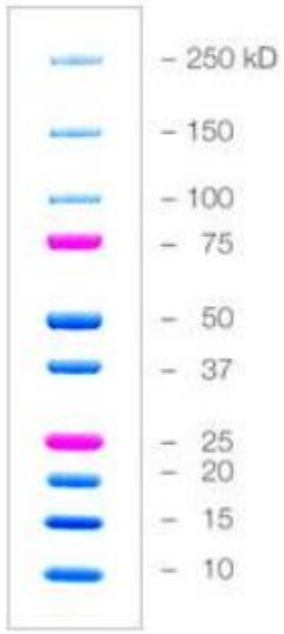


**Method used to capture the image** – The gel image was acquired used a Samsung A52. Blot images were acquired with a Vilber Fusion Fx (Vilber, France) imaging system using the PierceTM ECL Western kit.

Please refer to the manuscript for additional information, especially the paragraph “**Subtractive proteome analysis**” of the Methods section and the paragraph “**Identification of potential protein targets**” of the Results section.

#### Gel for gel excision (panel A)

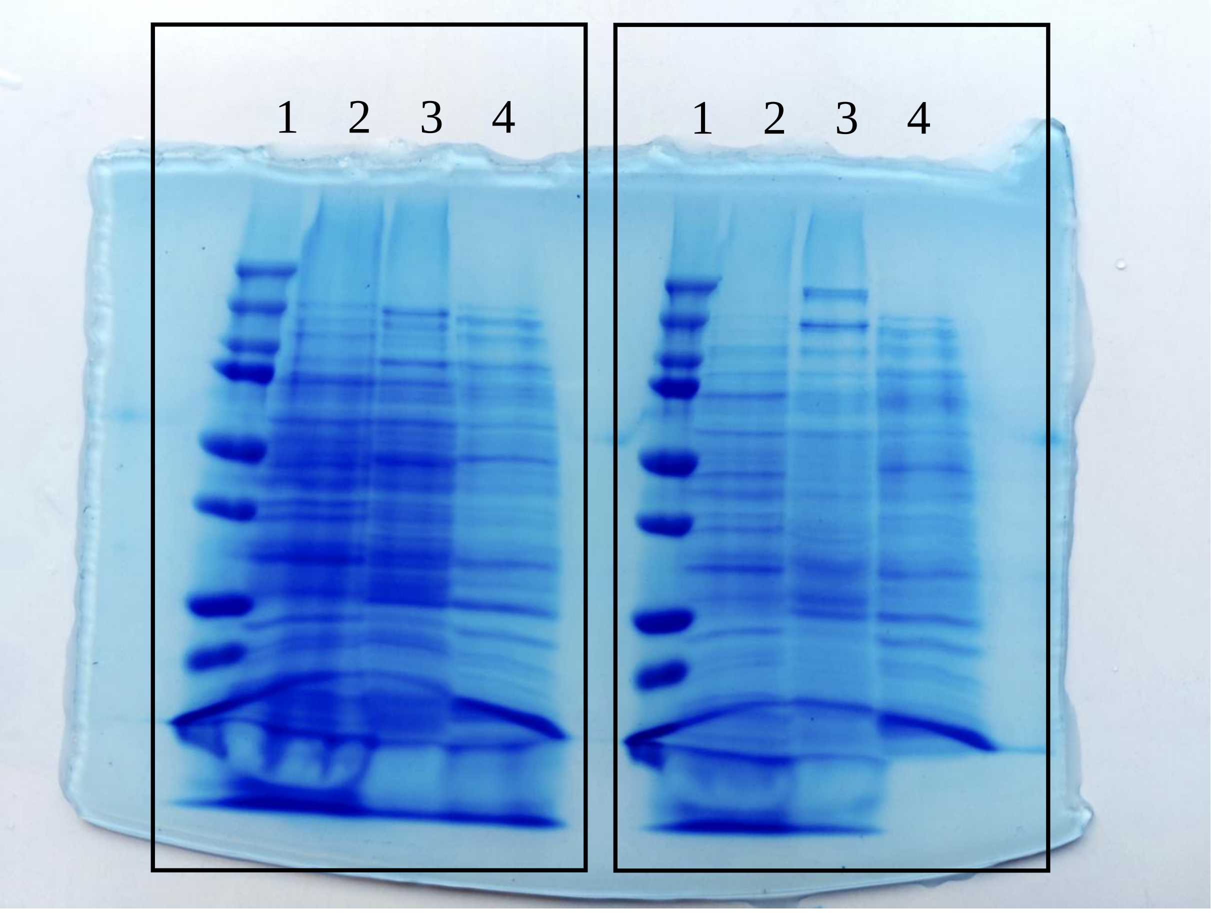


The left rectangle corresponds to the upper image in panel A: protein extractions from *S. aureus* MW2. The right rectangle corresponds to the lower image in panel A: protein extractions from *S. aureus* MN8.

#### **Blots incubated with non-depleted human sera (panel B)**

For MW2 (upper image in panel B of figure 3)

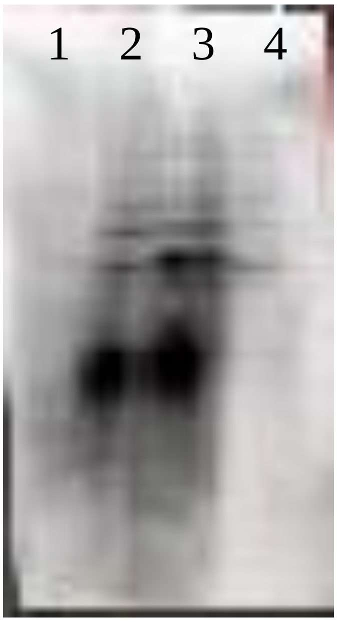


For MN8 (lower image in panel B of figure 3)

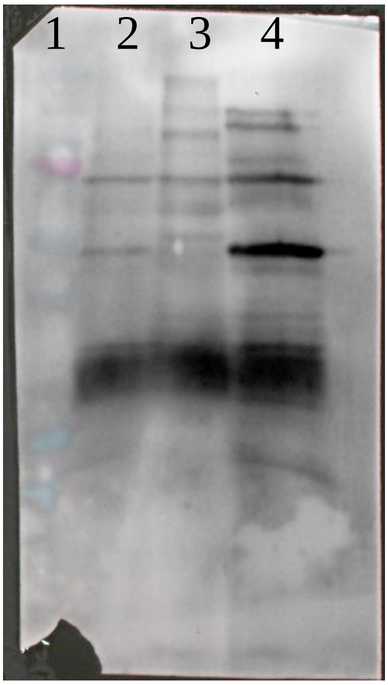

**Blots incubated with depleted human sera (panel C)**

For MW2 (upper image in panel C of figure 3)

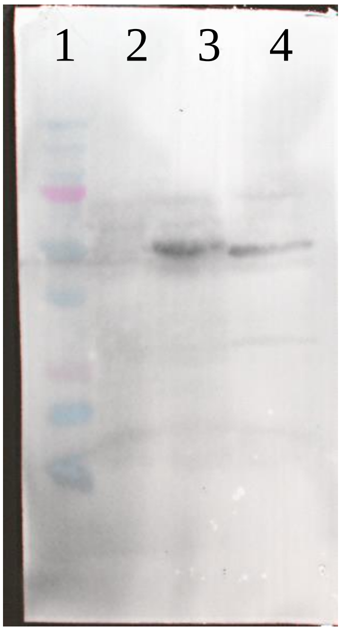


For MN8 (lower image in panel C of figure 3)

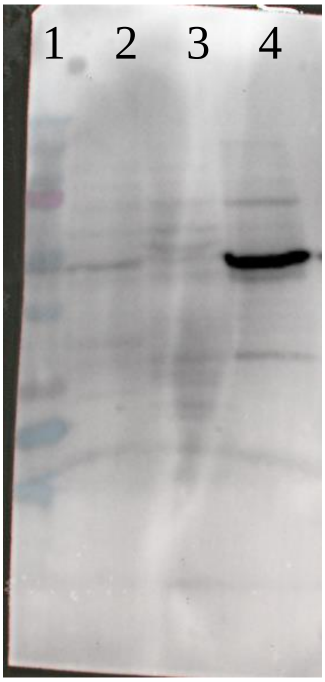
