## Supporting information for "Identification of cross-reactive vaccine antigen candidates in Gram-positive ESKAPE pathogens through subtractive proteome analysis using opsonic sera"

### Figure 2C

| | Human sera | | | Depleted Human Sera | | |
|---|---|---|---|---|---|---|
| **Dilution factor** | **$10^2$** | **$10^4$** | **$10^6$** | **$10^2$** | **$10^4$** | **$10^6$** |
| **Killing (%)** | 55,4 | 22,0 | 17,3 | 38,8 | 14,1 | 6,3 |
| **Killing (%)** | 63,3 | 27,0 | 21,8 | 38,8 | 11,1 | 8,9 |
| **Killing (%)** | 76,4 | 24,2 | 17,1 | 45,4 | 10,1 | 11,4 |
| **Killing (%)** | 54,1 | 21,5 | 12,6 | 45,4 | 5,2 | 3,8 |
| **Mean** | 62,3 | 23,7 | 17,2 | 42,1 | 10,1 | 7,6 |

### Figure 2D

| | Human sera | | | Depleted Human Sera | | |
|---|---|---|---|---|---|---|
| **Dilution factor** | **$10^2$** | **$10^4$** | **$10^6$** | **$10^2$** | **$10^4$** | **$10^6$** |
| **Killing (%)** | 50,6 | 29,2 | 0,1 | 30,4 | 17,8 | 2,0 |
| **Killing (%)** | 55,6 | 31,0 | 10,9 | 36,8 | 15,6 | 9,8 |
| **Killing (%)** | 39,1 | 23,9 | 3,5 | 35,2 | 24,4 | 1,1 |
| **Killing (%)** | 42,4 | 27,4 | 1,7 | 27,3 | 20,0 | 0,5 |
| **Mean** | 46,9 | 27,9 | 4,0 | 32,4 | 19,4 | 3,4 |

### Figure 5A

| | Pre-$AdcA_{au}$ | | | Anti-$AdcA_{au}$ | | |
|---|---|---|---|---|---|---|
| **Concentration µg/ml** | **0,5** | **0,25** | **0,125** | **0,5** | **0,25** | **0,125** |
| **$Abs_{405nm}$** | 0,093 | 0,086 | 0,082 | 4,000 | 2,887 | 1,732 |
| **$Abs_{405nm}$** | 0,094 | 0,088 | 0,081 | 3,999 | 3,340 | 1,985 |
| **$Abs_{405nm}$** | 0,091 | 0,087 | 0,083 | 4,000 | 3,111 | 1,857 |
| **$Abs_{405nm}$** | 0,095 | 0,087 | 0,080 | 4,000 | 3,117 | 1,861 |
| **Mean** | 0,094 | 0,087 | 0,082 | 4,000 | 3,114 | 1,859 |

### Figure 5B

| | Pre-AdcA$_{au}$ | | | Anti-AdcA$_{au}$ | | |
|---|---|---|---|---|---|---|
| **Concentration µg/ml** | **60** | **30** | **15** | **60** | **30** | **15** |
| **Abs$_{405nm}$** | 0,104 | 0,098 | 0,099 | 1,973 | 1,361 | 0,866 |
| **Abs$_{405nm}$** | 0,103 | 0,117 | 0,089 | 2,402 | 1,653 | 1,192 |
| **Abs$_{405nm}$** | 0,100 | 0,094 | 0,088 | 2,180 | 1,487 | 0,886 |
| **Abs$_{405nm}$** | 0,098 | 0,093 | 0,090 | 2,196 | 1,527 | 1,172 |
| **Mean** | 0,101 | 0,101 | 0,092 | 2,188 | 1,507 | 1,029 |

### Figure 6A

| | Pre-AdcA$_{au}$ | Anti-AdcA$_{au}$ | | | |
|---|---|---|---|---|---|
| **Dilution** | **40** | **40** | **80** | **160** | **320** |
| **Killing (%)** | 2,2 | 37,2 | 28,6 | 15,2 | 22,6 |
| **Killing (%)** | 4,2 | 28,2 | 38,0 | 26,2 | 29,0 |
| **Killing (%)** | 0,2 | 40,6 | 25,4 | 26,5 | 9,7 |
| **Killing (%)** | 7,6 | 43,3 | 22,5 | 36,4 | 19,4 |
| **Mean** | 3,6 | 37,3 | 28,6 | 26,1 | 20,2 |

### Figure 6B

| | Anti-AdcA$_{au}$ | | | | |
|---|---|---|---|---|---|
| **Inhibitor (µg/ml)** | **0** | **200** | **100** | **40** | **8** |
| **Killing (%)** | 50,3 | 7,8 | 27,5 | 34,1 | 38,2 |
| **Killing (%)** | 53,0 | 8,6 | 22,3 | 43,6 | 47,6 |
| **Killing (%)** | 48,9 | 4,6 | 33,5 | 34,1 | 53,0 |
| **Killing (%)** | 32,8 | 14,4 | 31,5 | 36,8 | 43,6 |
| **Mean** | 46,2 | 8,8 | 28,7 | 37,2 | 45,6 |

### Figure 7

| | *S. aureus Reynolds* 1:20 | | *E. faecium 11236/1* 1:20 | | *E. faecalis 12030* 1:50 | | *E. faecalis Type 2* 1:10 | |
|---|---|---|---|---|---|---|---|---|
| **Serum** | **Pre-AdcA$_{au}$** | **Anti-AdcA$_{au}$** | **Pre-AdcA$_{au}$** | **Anti-AdcA$_{au}$** | **Pre-AdcA$_{au}$** | **Anti-AdcA$_{au}$** | **Pre-AdcA$_{au}$** | **Anti-AdcA$_{au}$** |
| **Killing (%)** | 21,1 | 49,5 | 18,5 | 54,9 | 37,8 | 79,8 | 19,4 | 48,7 |
| **Killing (%)** | 26,3 | 48,6 | 21,5 | 66,9 | 35,0 | 78,3 | 3,3 | 48,7 |
| **Killing (%)** | 33,1 | 49,5 | 32,1 | 51,9 | 43,3 | 73,6 | 24,4 | 48,7 |
| **Killing (%)** | 22,9 | 49,2 | 16,5 | 51,9 | 35,0 | 82,9 | 7,3 | 55,9 |
| **Mean** | 25,9 | 49,2 | 22,2 | 56,4 | 37,8 | 78,7 | 13,6 | 50,5 |
